# Simulating Contact Using the Elastic Foundation Algorithm in OpenSim

**DOI:** 10.1101/319616

**Authors:** Michael W. Hast, Brett G. Hanson, Josh R. Baxter

## Abstract

Modeling joint contact is necessary to test many questions using simulation paradigms, but this portion of OpenSim is not well understood. The purpose of this study was to provide a guide for implementing a validated elastic foundation contact model in OpenSim. First, the load-displacement properties of a stainless steel ball bearing and ultra high molecular weight polyethylene (UHMWPE) slab were recorded during a controlled physical experiment. These geometries were imported and into OpenSim and contact mechanics were modeled with the on-board elastic foundation algorithm. Particle swarm optimization was performed to determine the elastic foundation model stiffness (2.14×10^11^ ± 6.81×10^9^ N/m) and dissipation constants (0.999 ± 0.003). Estimations of contact forces compared favorably with blinded experimental data (root mean square error: 87.58 ± 1.57 N). Last, total knee replacement geometry was used to perform a sensitivity analysis of material stiffness and mesh density with regard to penetration depth and computational time. These simulations demonstrated that material stiffnesses between 10^11^ and 10^12^ N/m resulted in realistic penetrations (< 0.15mm) when subjected to 981N loads. Material stiffnesses between 10^13^ and 10^15^ N/m increased computation time by factors of 12–23. This study shows the utility of performing a simple physical experiment to tune model parameters when physical components of orthopaedic implants are not available to the researcher. It also demonstrates the efficacy of employing the on-board elastic foundation algorithm to create realistic simulations of contact between orthopaedic implants.

## Introduction

Predicting articular joint function and loading continues to be an important research topic in orthopaedics. Effective clinical treatment and design of orthopaedic implants requires a thorough understanding of joint loads throughout dynamic activities of daily living. Computer simulation has become a widely used method for the determination of joint contact forces during dynamic tasks, as it is not subject to the same constraints that accompany physical experimental investigations. Modeling joint contact using an elastic foundation (EF) paradigm is a commonly utilized approach because of its cheap computational cost, a desirable characteristic for integrating joint contact into muscle-driven simulations (Kim et al., 2009; Lenhart et al., 2015; Lin and Fregly, 2010; Schmitz and Piovesan, 2016; Shelburne et al., 2006; Taylor et al., 2004).

OpenSim (Delp et al., 2007) is a widely used and freely available musculoskeletal modeling software package that has an onboard EF contact algorithm. Based on the history of forum posts and a dearth of publications using this paradigm, this Opensim feature is not well understood by many users (Dunne et al., 2013, 2017a, 2017b). Perhaps for this reason, the on-board algorithm is seldom used for the purposes of estimating joint forces. Although the mathematical concept behind the EF algorithm is outlined in several publications (Sherman et al., 2011; Uchida et al., 2015), details regarding validation and day-to-day use remain confusing for end users. For example, the material stiffness constant is described with the following equation:

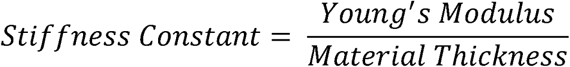

It is unclear if other variables within the model (dissipation, mesh density, geometric complexity) affect accuracy and computation time, two important metrics of performance in computational modeling.

The purpose of the present work was to provide end users with a better understanding of the on-board EF algorithm in OpenSim, so that estimations of joint contact forces can be made readily, with accuracy, and with consistency. To do this, we developed a simple experiment and concomitant OpenSim models that provide guidelines for determining EF input parameters. For simplicity, a validation experiment was first performed to investigate the estimation of contact forces between a sphere and a plate. To demonstrate a more complex and biomechanically relevant simulation, contact of a total knee arthroplasty (TKA) was simulated to test the sensitivity of simulation performance to small changes to model parameters.

## Methods

### Tuning and Testing Contact Parameters: Sphere-on-Plate

Elastic foundation parameters for simple model were established using a physical experiment. A tightly toleranced 5.08 cm diameter 316L stainless steel ball bearing and a 15.24 × 7.62 × 0.95 cm thick slab of ultra high molecular weight polyethylene (UHMWPE) underwent mechanical testing. The UHMWPE rested freely on the bed of the test frame (Electroforce 3330, TA Instruments, New Castle, DE) and the sphere was placed on the slab, directly under the actuator (Figure 1A). To simulate dynamic loading, the actuator imparted loads between 0 – 750 N at 1 Hz for 50 cycles while force-displacement data were collected at 100 Hz. This protocol was repeated 10 times, taking care to use unblemished areas of the deformable plastic for each trial.

**Figure 1.**
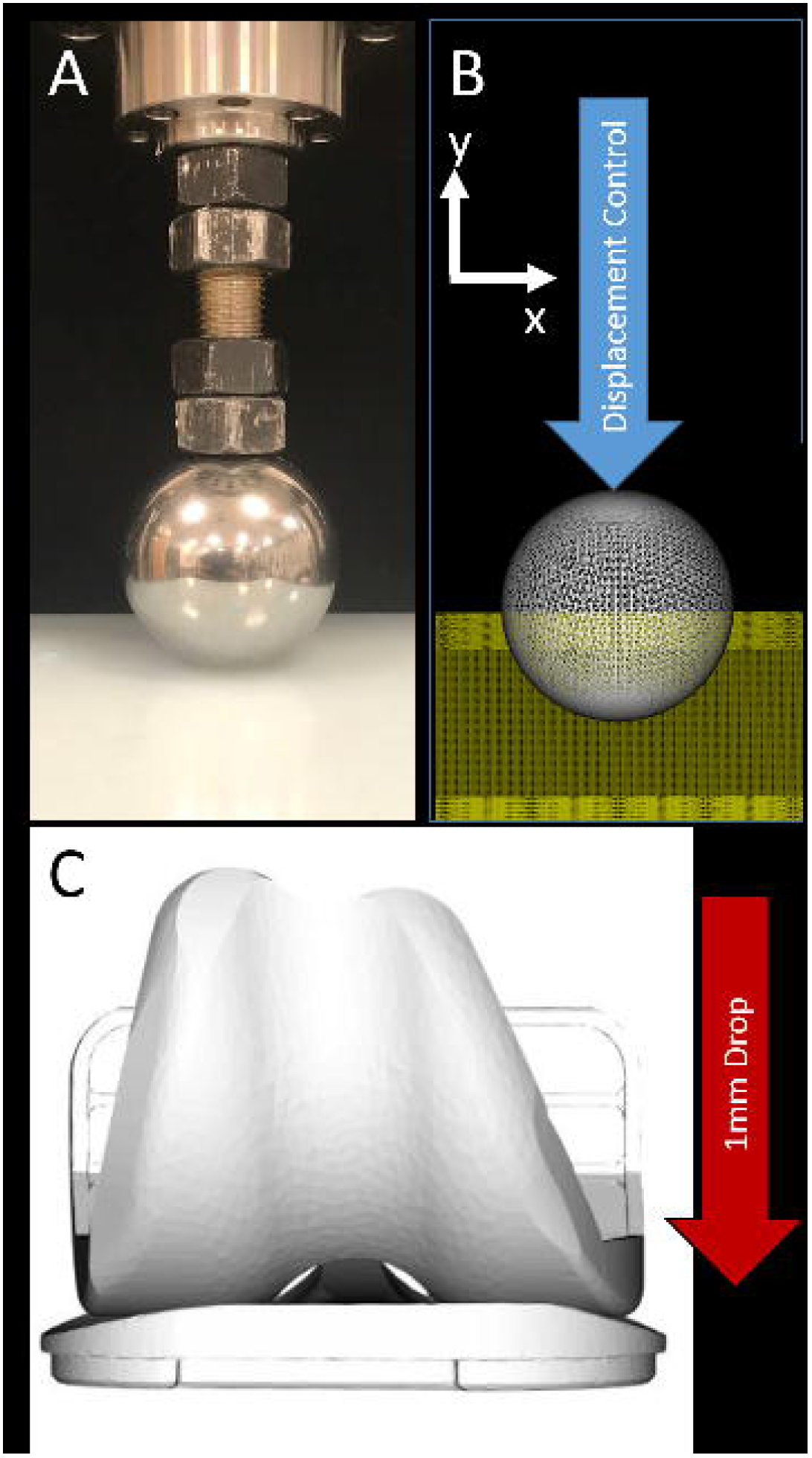
(A) A photograph showing the experimental setup involving a stainless steel sphere in contact with a UHMWPE plate. The actuator of the test frame imposed cyclic loads of 750N upon the sphere. (B) A screenshot from the computational model of the physical experiment. The measured displacements of the sphere during the physical experiment were replicated, so that the stiffness and dissipation constants could be tuned. (C) “Drop and settle” simulations containing TKA components, were performed to examine the relationships between stiffness, mesh density, and computation time.

Computer generated three dimensional (3D) renderings of the sphere and plate (Solidworks 2017, Dassault Systèmes, Waltham, MA) were imported into OpenSim (version 3.3, Appendix A) and defined as an EF contact model. Prior to being imported into OpenSim, both 3D bodies were converted into watertight stereolithography (STL) files in American Standard Code for Information Exchange (ASCII) format. Because EF models predict forces based upon intersections of triangular faces of the 3D bodies, the geometries were resampled to ensure the results from this study could be transferred to more complex geometries. Specifically, the sphere was made up of 5852 faces and oriented such that the pole was in contact with the plate. Rather than representing the rectangular plate with 12 triangular faces, the mesh geometry was subdivided using open-source mesh editing software (Meshlab, (Cignoni et al., 2008)) to have 30208 faces (Figure 1B).

Stiffness and dissipation constants in the EF model were tuned by using an optimization approach in the following manner. Based on pilot testing, the first 30 loading cycles of each 50 cycle trial exhibited substantial hysteresis. These cycles were omitted to allow for preconditioning the UHMWPE slab. The force displacement profiles of the 31^st^–40^th^ trials were used to tune the computational model by simulating prescribed motions in OpenSim that exactly replicated the displacements measured during the physical experiment. Contact forces between the two bodies were calculated in OpenSim with the onboard Force Reporter algorithm. The root mean square error (RMSE) of the force versus time curve for the physical experiment and simulation was used as an objective function. For each experimental trial, the values of stiffness and dissipation were optimized using a hybrid “particleswarm/fmincon” function (MATLAB, The Mathworks, Natick, MA) until the RMSE was minimized. Parameter values from all trials were averaged to create optimized constants for stiffness and dissipation. Finally, using the averaged constants, the displacements of the 40^th^–49^th^ cycles of the experimental trials were simulated and the resulting contact forces were estimated. The 50^th^ cycle was omitted because completion of the last cycle in physical experiment was not consistent across trials. To assess tuning accuracy, the newly estimated force versus time curves were compared to their measured counterparts by calculating the RMSE of the two curves.

### Simulating contact mechanics of total knee arthroplasty

Tibiofemoral contact was modeled using a forward dynamic simulation to quantify the relationships between computation time, implant penetration, material stiffness, and mesh density. Computer aided designs of a cruciate-retaining total knee replacement (eTibia (Fregly et al., 2012)) were imported into OpenSim and contact was modeled as an elastic foundation element. ‘Drop and settle’ simulations (Figure 1C) were performed over a series of model configurations to test the effects of tibial insert stiffness (10^9^ to 10^15^ N/m) mesh coarseness (100, 500, 1,000, 2,500, and 5,000 faces), and computational time. Different amounts of weight bearing were also simulated by changing the mass of the femoral component (1, 10, and 100kg). Center of mass position was adjusted to eliminate rotation of the femoral component during simulations. Component penetration was calculated as the vertical change in position of the femoral component center of mass from the point of initial component contact to the final settling position. In total, 105 simulations were performed on a personal computer (Intel Core i5-6500, 3.20GHz, 8GB RAM).

## Results and Discussion

Stiffness and dissipation constants accurately predicted experimental cyclic loading mechanics (Figure 2). The tuning optimization yielded average stiffness and dissipation constants of 2.14×10^11^ ± 6.81×10^9^ and 0.999 ± 0.003, respectively. The average stiffness constant found in the current study falls just outside of the range of stiffness constants that are determined when dividing the modulus of UHMWPE (780 – 990 MPa (Kurtz, 2004)) by the thickness of 0.95 cm (8.2×10^10^−1.0×10^11^ N/m).

**Figure 2.**
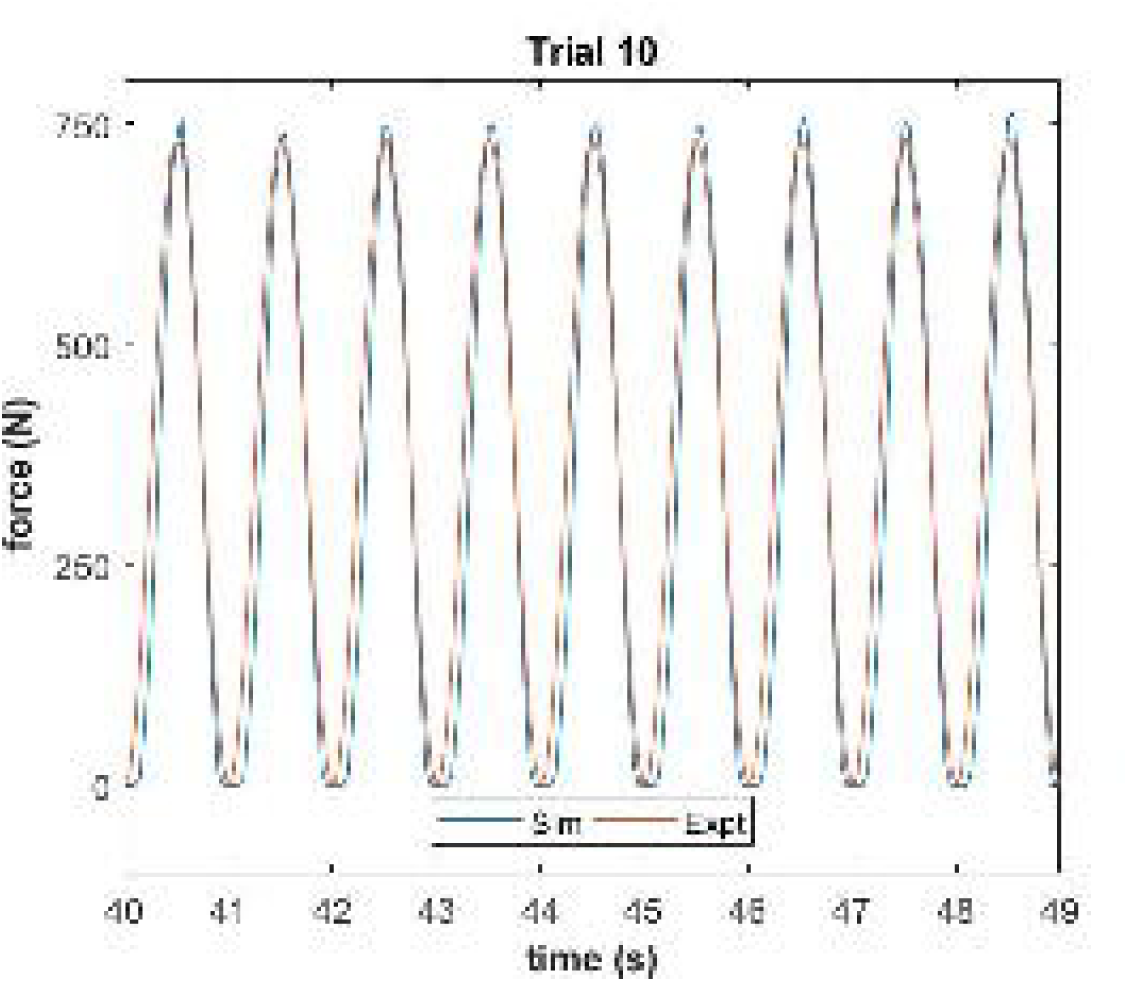
A plot showing the experimental force versus time plot (orange) in comparison to the estimated forces (blue). Simulated forces were estimated based upon the optimization results for stiffness and dissipation constants. This plot represents only one trial. Plots for all trials can be found in Appendix B.

When comparing the experimental force versus time curves to the simulated ones, the average root mean squared error (RMSE) was 87.58 ± 1.57 N. Because some differences were due to a slight phase lag in the simulations, the estimated and measured peak loads for all trials were also compared with a RMSE technique. In this case, the average RMSE value was 32.45 ± 23.54 N. A table of results from all trials can be found in Appendix B.

Simulated tibiofemoral penetration was below the *a priori* threshold of 0.15 mm (Muratoglu et al., 2003) with the stiffness set at 10^11^ and 10^12^ N/m and the contact mesh having at least 1000 faces (Figure 3). Lower stiffness values (10^9^ and 10^10^ N/m) and higher stiffness values (10^13^ to 10^15^ N/m) resulted in non-physiologic penetrations of greater than 0.5 mm and component chattering, respectively. Computation time increased as a function of both material stiffness and mesh density; however, overly stiff (10^14^–10^15^ N/m) increased by a factor of 12 – 23 times longer than simulations with 10^11^ N/m.

**Figure 3.**
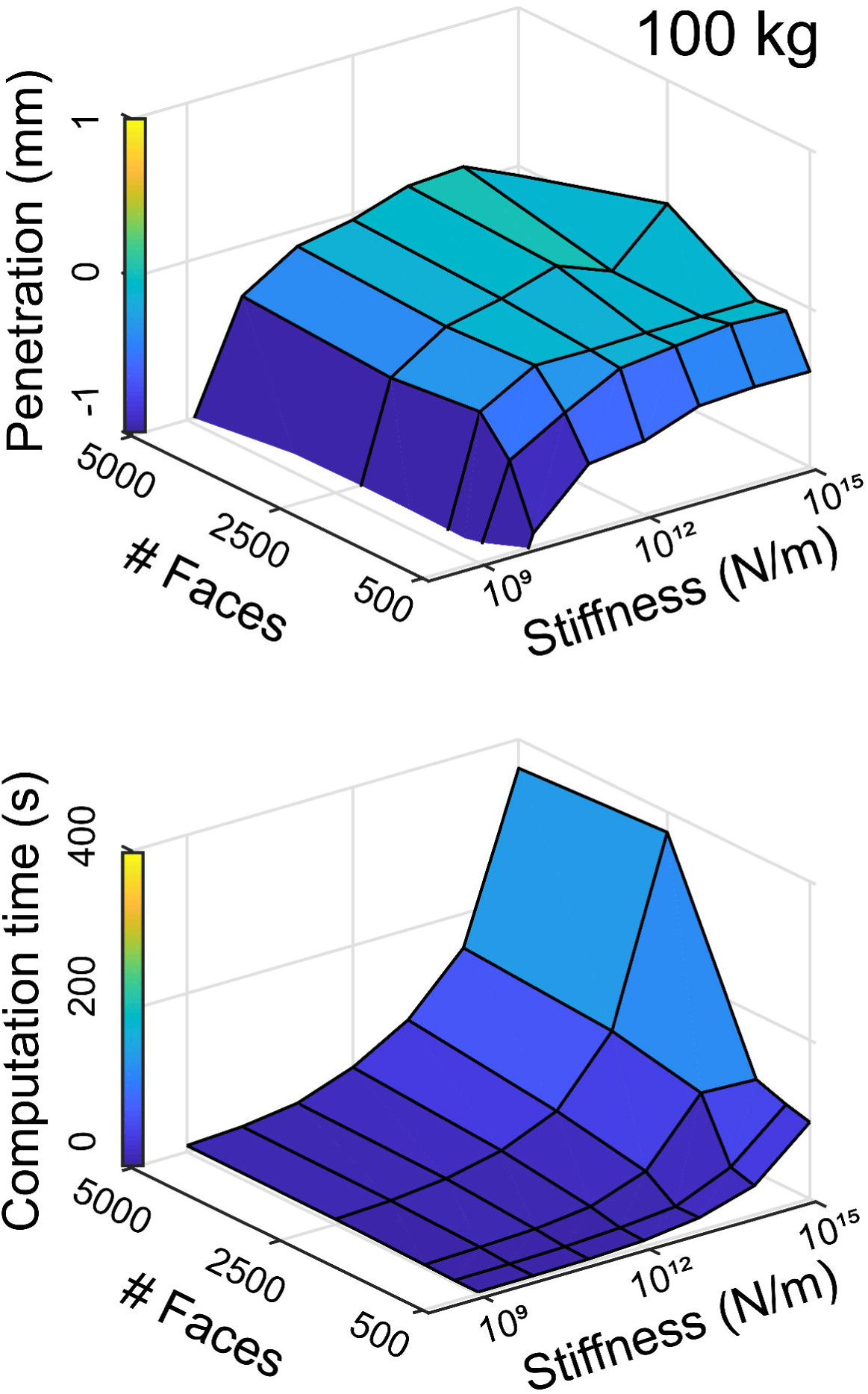
Component penetration (top row) and computation time (bottom row) were more sensitive to material stiffness than the mesh density (# faces). Low and stiffness constants resulted in excess penetration (negative values) and increased computation time, respectively.

Contact mechanics can be easily simulated using an EF paradigm when the appropriate model parameters are selected. Component penetration is dependent on the stiffness coefficient, where increased stiffness drastically increases computation time and creates chattering behaviors between the contacting bodies, which is an artifact of increased joint reaction loads (Appendix C). Mesh size also plays an important role in overall simulation performance. Course meshes resulted in increased component penetration and fine meshes increased computation time unnecessarily.

This experiment has several limitations. In the sphere on plate model, the maximum load of 750 N was chosen because it represents approximately one body weight of a 50^th^ percentile male; however, the small contact patch between the plate and sphere resulted in slightly higher maximum contact pressures (24.97 ± 4.04 MPa Appendix D) than reported in TKA (~19 MPa) (Kwon et al., 2014). Although the current experiment represents a worst-case scenario, future experiments may consider utilizing more conforming geometries to better represent realistic stresses. The parameters of static, dynamic, and viscous friction were all assumed to be negligent and therefore were set to zero (Hast and Piazza, 2013; Thompson et al., 2011). This may not be the case for more complex motions, and the parameters could readily be added to the optimization routine. Finally, the TKA experiment was not validated with physical experiments. Such an effort would require physical TKA implants and matching CAD geometries, which were not available for this experiment.

## Conclusions

OpenSim provides an EF algorithm that is freely available, computationally light, and potentially powerful, but the tool is vastly underutilized because it is poorly understood. This experiment represents the first published work that has outlined a rigorous experimental approach to determine constants that will accurately predict forces using the OpenSim EF contact algorithm. For the purposes of simplicity and practicality, a simple stainless steel sphere and UHMWPE plate were used in the physical experiment, which translated favorably into a virtual model of a TKA. When considering other materials or other joints of the body, the same overall approach can be straightforwardly adapted to make reasonable estimations joint contact forces.

## Acknowledgment

The authors would like to thank the Biedermann family, who provided funding for this study.

